# Mitochondrial one-carbon enzyme MTHFD1L sustains stemness and metastatic progression in breast cancer

**DOI:** 10.64898/2026.06.18.732629

**Authors:** Hirokazu Kusunoki, Tsunaki Hongu, Tatsunori Nishimura, Kazuki Ogawa, Jin Lee, Yasuto Takeuchi, Koji Okamoto, Tomoyoshi Soga, Noriko Gotoh

## Abstract

One-carbon (1C) metabolism is frequently upregulated in cancer to support anabolic growth and nucleotide biosynthesis. However, the contribution of mitochondrial 1C metabolism to cancer stemness and metastatic progression remains incompletely understood. Here, we identify methylenetetrahydrofolate dehydrogenase 1-like (MTHFD1L), a mitochondrial 1C metabolic enzyme, as a critical regulator of breast cancer stemness, tumor initiation, and lung metastasis. Genetic depletion of *MTHFD1L* in triple-negative breast cancer (TNBC) cell lines and patient-derived breast cancer models markedly impaired proliferation, sphere formation, tumorigenicity, and lung colonization. Metabolomic profiling revealed extensive metabolic rewiring following MTHFD1L loss, characterized by accumulation of the purine biosynthetic intermediates SAICAR and AICAR together with perturbations in glycolytic and pentose phosphate pathways. Importantly, suppression of lung metastasis was accompanied by reduced expression of the stemness-associated transcription factor SOX2 and decreased proliferative activity in metastatic lesions. Collectively, our findings establish MTHFD1L as a key metabolic dependency linking mitochondrial 1C metabolism to stem-like properties and metastatic progression in breast cancer, and highlight MTHFD1L as a promising therapeutic target in metastatic TNBC.

## 1. Introduction

Metastatic progression remains the leading cause of cancer-related mortality worldwide^1^. Despite substantial advances in targeted therapies and immunotherapy, disseminated cancer cells frequently survive therapeutic intervention and establish secondary lesions in distant organs. Increasing evidence suggests that metastatic competence is driven by highly plastic cancer cell populations capable of adapting to diverse microenvironmental and therapeutic stresses^2^. Understanding the mechanisms that support this adaptive capacity is therefore essential for developing more effective therapies against metastatic disease.

Breast cancer is the most frequently diagnosed malignancy in women and remains a major cause of cancer-related death^3^. Among its molecular subtypes, triple-negative breast cancer (TNBC), characterized by the absence of estrogen receptor (ER), progesterone receptor (PR), and human epidermal growth factor receptor 2 (HER2), is associated with particularly poor clinical outcomes because of its aggressive behavior, high metastatic potential, and lack of effective targeted therapies^4^. Metastatic dissemination exposes cancer cells to profound metabolic and environmental stresses, requiring dynamic adaptation to support survival and colonization of distant organs^2^. Notably, stem-like cancer cell populations have been implicated in tumor initiation, therapeutic resistance, metastatic progression, and disease recurrence, suggesting that specialized metabolic programs may contribute to the maintenance of these aggressive cellular states ^5,6^.

Metabolic reprogramming is now recognized as a hallmark of cancer^7^. Among the metabolic pathways implicated in tumor progression, folate-mediated one-carbon (1C) metabolism occupies a central position by providing 1C units required for nucleotide biosynthesis and redox homeostasis^8–11^. One-carbon metabolism has long been therapeutically targeted in cancer, exemplified by the clinical use of methotrexate (MTX). However, global inhibition of folate metabolism often causes substantial toxicity in normal proliferating tissues, highlighting the need for more selective therapeutic approaches.

In mammalian cells, 1C metabolism is compartmentalized between the cytosol and mitochondria^12^. Within mitochondria, serine-derived 1C units are converted into formate through sequential enzymatic reactions catalyzed by serine hydroxymethyltransferase 2 (SHMT2), methylenetetrahydrofolate dehydrogenase 2 (MTHFD2), and methylenetetrahydrofolate dehydrogenase 1-like (MTHFD1L) (Figure 1a). Mitochondria-derived formate subsequently supports cytosolic purine and pyrimidine biosynthesis.

**Fig. 1.**
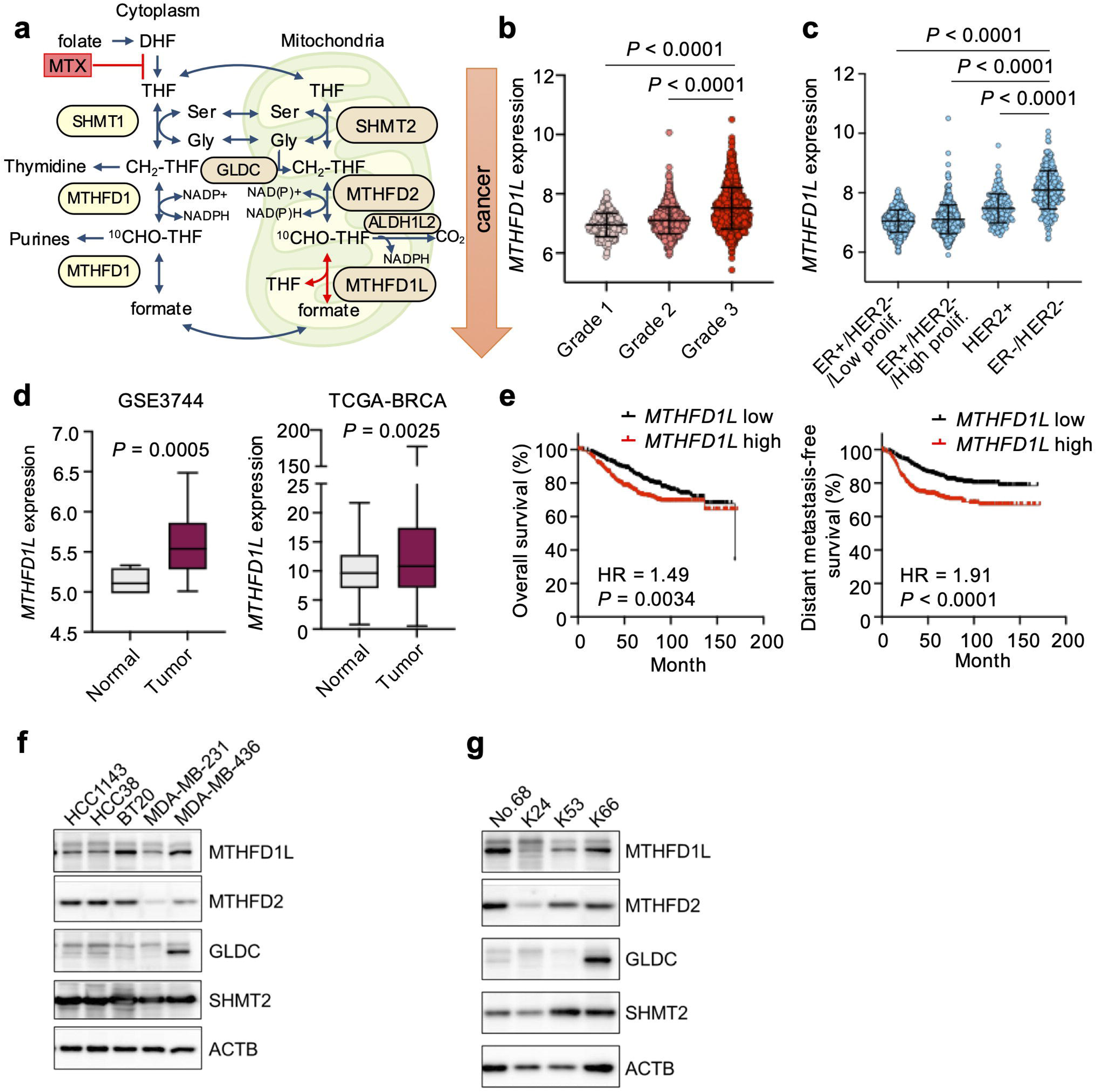
MTHFD1L expression is associated with breast cancer progression and poor prognosis. (a) Schematic overview of folate-mediated one-carbon (1C) metabolism. The reaction catalyzed by methylenetetrahydrofolate dehydrogenase 1-like (MTHFD1L) is highlighted in red. Methotrexate (MTX) blocks the reaction from dihydrofolate (DHF) to tetrahydrofolate (THF). SHMT1/2 (Serine hydroxymethyltransferase 1 and 2), MTHFD1/2 (methylenetetrahydrofolate dehydrogenase 1 and 2), GLDC (Glycine decarboxylase), and ALDH1L2 (aldehyde dehydrogenase 1 family member 2). (b) *MTHFD1L* expression according to tumor grade in breast cancer. (c) *MTHFD1L* expression according to breast cancer molecular subtype. ER, estrogen receptor; HER2, human epidermal growth factor receptor 2; prolif, proliferation. P values in (b) and (c) were determined by one-way ANOVA with Tukey’s multiple comparisons test. Data are shown as mean ± SD. (d) *MTHFD1L* expression in normal breast tissues and breast tumor tissues analyzed using the GSE3744 dataset (left) and TCGA-BRCA dataset (right). Boxes indicate the median and interquartile range, and whiskers indicate the minimum and maximum values. P values were determined by two-tailed Mann–Whitney test. (e) Kaplan–Meier analysis of overall survival and distant metastasis-free survival in breast cancer patients using the Kaplan–Meier Plotter dataset. Patients were stratified according to median *MTHFD1L* expression. n = 943 patients for overall survival and n = 958 patients for distant metastasis-free survival. P values were determined by log-rank test. HR, hazard ratio. (f,g) Expression of one-carbon metabolic enzymes in breast cancer cell lines (f) and patient-derived breast cancer cells (g).

Notably, mitochondrial 1C metabolic enzymes are frequently upregulated in cancer and have emerged as attractive therapeutic targets. Compared with conventional antifolate therapies, selective inhibition of mitochondrial 1C metabolism may provide a broader therapeutic window while preserving normal tissue function ^9–11^. Indeed, we previously demonstrated that mammary epithelial cell-specific deletion of MTHFD2 does not impair pregnancy-induced mammary gland development, suggesting that normal cell growth is largely dispensable for mitochondrial 1C metabolism^13^. These findings raise the possibility that cancer cells may exhibit a greater dependence on mitochondrial 1C metabolism than normal tissues, thereby creating a potential therapeutic vulnerability.

Among mitochondrial 1C enzymes, SHMT2 and MTHFD2 have been extensively investigated and have been implicated in tumor growth, stress adaptation, and stem-like properties^9^. We previously demonstrated that MTHFD2 contributes to maintenance of stemness in lung cancer^14^. In contrast, the functional role of MTHFD1L in breast cancer progression remains incompletely understood. Although MTHFD1L catalyzes the final step of mitochondrial formate production and occupies a central position in mitochondrial 1C metabolism, its contribution to tumor initiation, metastatic progression, and the maintenance of stem-like cancer cell states remains poorly understood.

In the present study, we investigated the role of MTHFD1L in TNBC using breast cancer cell lines, patient-derived cancer cells and patient-derived models. We demonstrate that MTHFD1L is required for maintenance of stem-like properties, tumor initiation, and lung colonization. Furthermore, metabolomic analyses reveal extensive metabolic rewiring following *MTHFD1L* depletion, characterized by accumulation of nucleotide biosynthetic intermediates and disruption of interconnected metabolic pathways. Our findings establish MTHFD1L as a critical metabolic dependency in aggressive breast cancer and support therapeutic targeting of mitochondrial 1C metabolism as a strategy to suppress metastatic progression.

## 2. Materials and Methods

### 2.1. Lung colonization assay

Female NOD/SCID/IL2Rγ^null^ (NSG) mice aged 6–12 weeks were maintained under pathogen-free conditions according to institutional guidelines. Animal studies were approved by the Animal Research Committee of Kanazawa University.

MDA-MB-231 or No.68 cells (2 x 10^5^)^15,16^ expressing the triple-reporter fusion gene (TGL) reporter consisting of herpes simplex virus thymidine kinase (HSV1-TK), green fluorescent protein (GFP), and firefly luciferase (Fluc), were suspended in phosphate-buffered saline (PBS) and injected intravenously through the tail vein. Lung colonization was monitored by bioluminescence imaging using an IVIS Lumina LT imaging system (PerkinElmer Waltham, MA). Mice were injected intraperitoneally with D-luciferin (Wako, Osaka, Japan) (150 mg/kg) prior to imaging. Bioluminescence signals were quantified using Living Image software.

### 2.2. Statistical analysis

Statistical analyses were performed using GraphPad Prism 10 unless otherwise indicated. Data are presented as mean ± SD or mean ± SEM as specified in the figure legends. Comparisons between two groups were performed using Student’s t-test, Welch’s t-test, or Mann–Whitney U test as appropriate. Comparisons among multiple groups were performed using one-way or two-way ANOVA followed by appropriate multiple-comparison tests.

For RNA-seq analyses, differential gene expression was determined using DESeq2. For Gene set enrichment analysis (GSEA), significance was defined as nominal P < 0.05 and false discovery rate (FDR) < 0.25. Extreme limiting dilution analyses were performed using ELDA software. P values < 0.05 were considered statistically significant.

## 3. Results

### 3.1. MTHFD1L is associated with poor prognosis and promotes growth and stem-like properties of TNBC cells

To investigate the clinical relevance of MTHFD1L in breast cancer, we analyzed publicly available transcriptomic datasets, including the METABRIC cohort^17^. *MTHFD1L* expression was elevated in high-grade breast cancers and was particularly enriched in TNBC compared with other molecular subtypes (Figure 1b,c). Furthermore, *MTHFD1L* expression was significantly higher in tumor tissues than in normal breast tissues and was associated with poor clinical outcomes, including reduced overall survival and distant metastasis-free survival (Figure 1d,e) ^18^.

MTHFD1L, together with other mitochondrial 1C metabolic enzymes, including MTHFD2 and SHMT2, was broadly expressed in TNBC cell lines and patient-derived breast cancer cells (Figure 1f,g), whereas expression of another mitochondrial enzyme, GLDC, was more restricted.

To determine the functional contribution of MTHFD1L to breast cancer cell growth, we first depleted *MTHFD1L* using lentiviral shRNAs. *MTHFD1L* knockdown significantly impaired proliferation of breast cancer cells (Figure 2a,b). We next established a DOX-inducible CRISPR-Cas9 knockout (KO) system. DOX-induced deletion of *MTHFD1L* significantly reduced the growth of both patient-derived breast cancer cells (Figure 2c,d) and TNBC cell lines (Figure 2e–h). Importantly, supplementation with formate, a downstream product of mitochondrial 1C metabolism, substantially rescued the growth defect caused by *MTHFD1L* deletion (Figure 2f,h), confirming that the observed phenotype resulted from disruption of mitochondrial 1C metabolism.

**Fig. 2.**
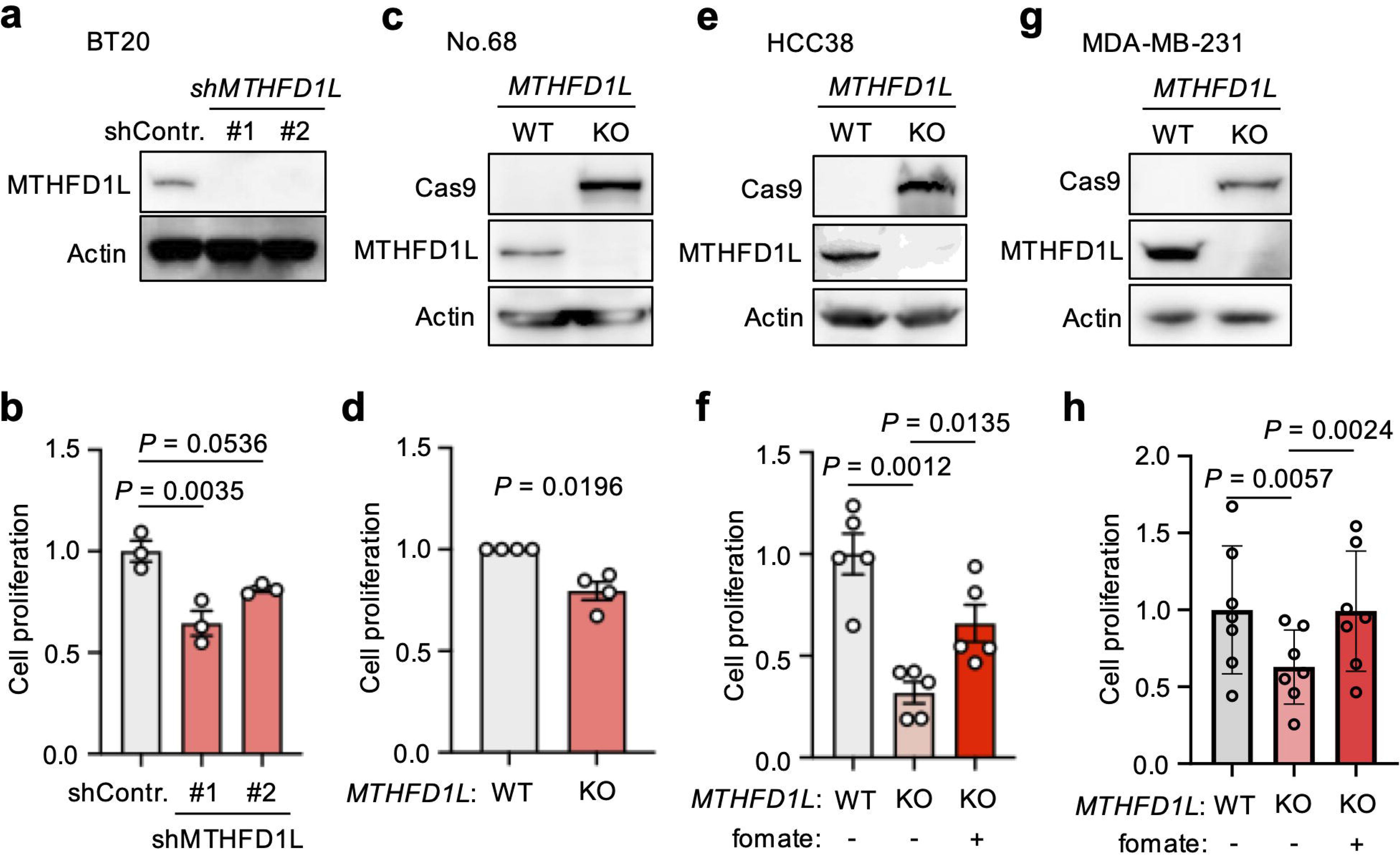
MTHFD1L promotes breast cancer cell growth in vitro. (a,c,e,g) Depletion of *MTHFD1L* in breast cancer cells. shRNA-mediated knockdown of *MTHFD1L* in BT20 cells (a) and CRISPR/Cas9-mediated doxycycline (DOX)-inducible knockout (KO) of *MTHFD1L* in No.68 patient-derived breast cancer cells (c), HCC38 cells (e), and MDA-MB-231 cells (g) are shown. WT, wild type. (b,d,f,h) Effects of *MTHFD1L* depletion on proliferation of BT20 cells (b), No.68 cells (d), HCC38 cells (f), and MDA-MB-231 cells (h). Data are shown as mean ± SEM from 3–6 independent experiments. P values were determined by one-way ANOVA with Dunnett’s multiple comparisons test in (b), (f), and (h), and by two-tailed paired t-test in (d).

Because sphere formation reflects stem-like properties and self-renewal capacity^15, 16^, we next examined whether MTHFD1L contributes to stemness. *MTHFD1L* knockdown significantly reduced sphere formation in both established breast cancer cell lines and patient-derived breast cancer cells (Figure 3a,b). Consistently, extreme limiting dilution analysis (ELDA) demonstrated a significant reduction in sphere-forming frequency following MTHFD1L depletion (Figure 3c–e). Collectively, these findings indicate that MTHFD1L promotes both proliferative capacity and stem-like properties in TNBC cells.

**Fig. 3.**
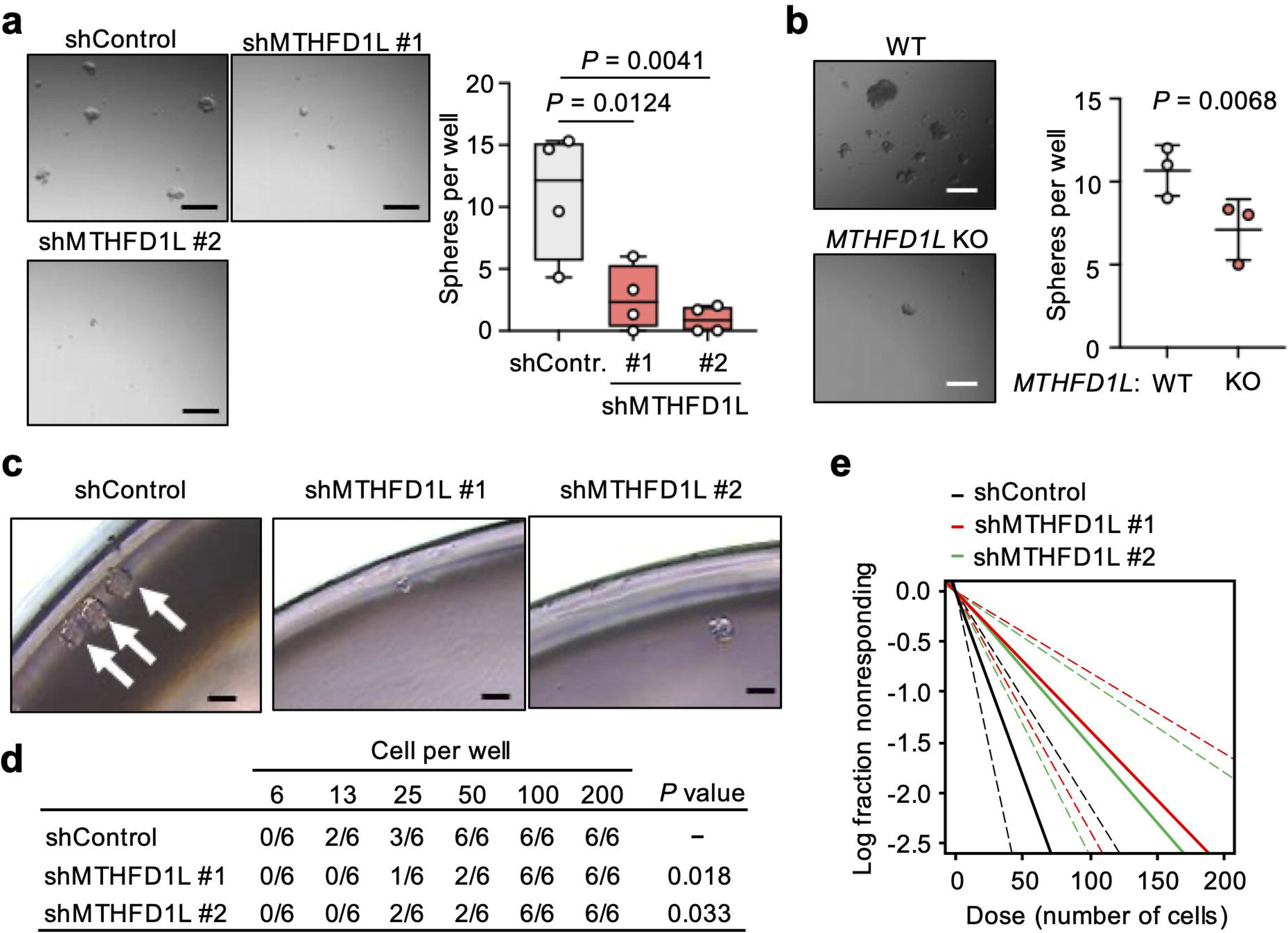
MTHFD1L supports sphere-forming ability in breast cancer cells. (a) Effects of *MTHFD1L* knockdown on sphere formation in BT20 breast cancer cells. Representative images of spheres are shown on the left, and quantification of the number of spheres per well is shown on the right. Scale bar, 200 µm. (b) Sphere formation of WT and *MTHFD1L*-KO No.68 cells. Representative images are shown on the left, and quantification of the number of spheres per well is shown on the right. Scale bar, 200 µm. P values were determined by one-way ANOVA with Dunnett’s multiple comparisons test from four independent experiments in (a), and by two-tailed paired t-test from three independent experiments in (b). (c–e) Extreme limiting dilution assay (ELDA) of BT20 cells following *MTHFD1L* depletion. Representative images (c) and sphere-forming frequency (d,e) are shown. Spheres larger than 75 µm after 5 days of incubation were counted. Arrows in (c) indicate spheres. P values were determined by goodness-of-fit test.

### 3.2. MTHFD1L depletion induces extensive metabolic rewiring

To investigate the metabolic consequences of MTHFD1L loss, we performed comprehensive metabolomic profiling and RNA sequencing using wild-type and MTHFD1L-KO HCC38 cells.

Pathway enrichment analysis revealed significant alterations in multiple metabolic pathways, including glycine, serine and threonine metabolism, pyrimidine metabolism, purine metabolism, and one carbon pool by folate (green letters in Figure 4a). Consistent with inhibition of mitochondrial 1C flux, serine accumulated significantly in MTHFD1L-KO cells, accompanied by an increase in the NADPH/NADP+ ratio (Figure 4b,c). Notably, RNA-seq analysis revealed little changes in the expression of mitochondrial or cytosolic 1C metabolic enzymes (Figure S1a), suggesting that the observed metabolic alterations primarily reflect changes in metabolic flux rather than transcriptional regulation.

**Fig. 4.**
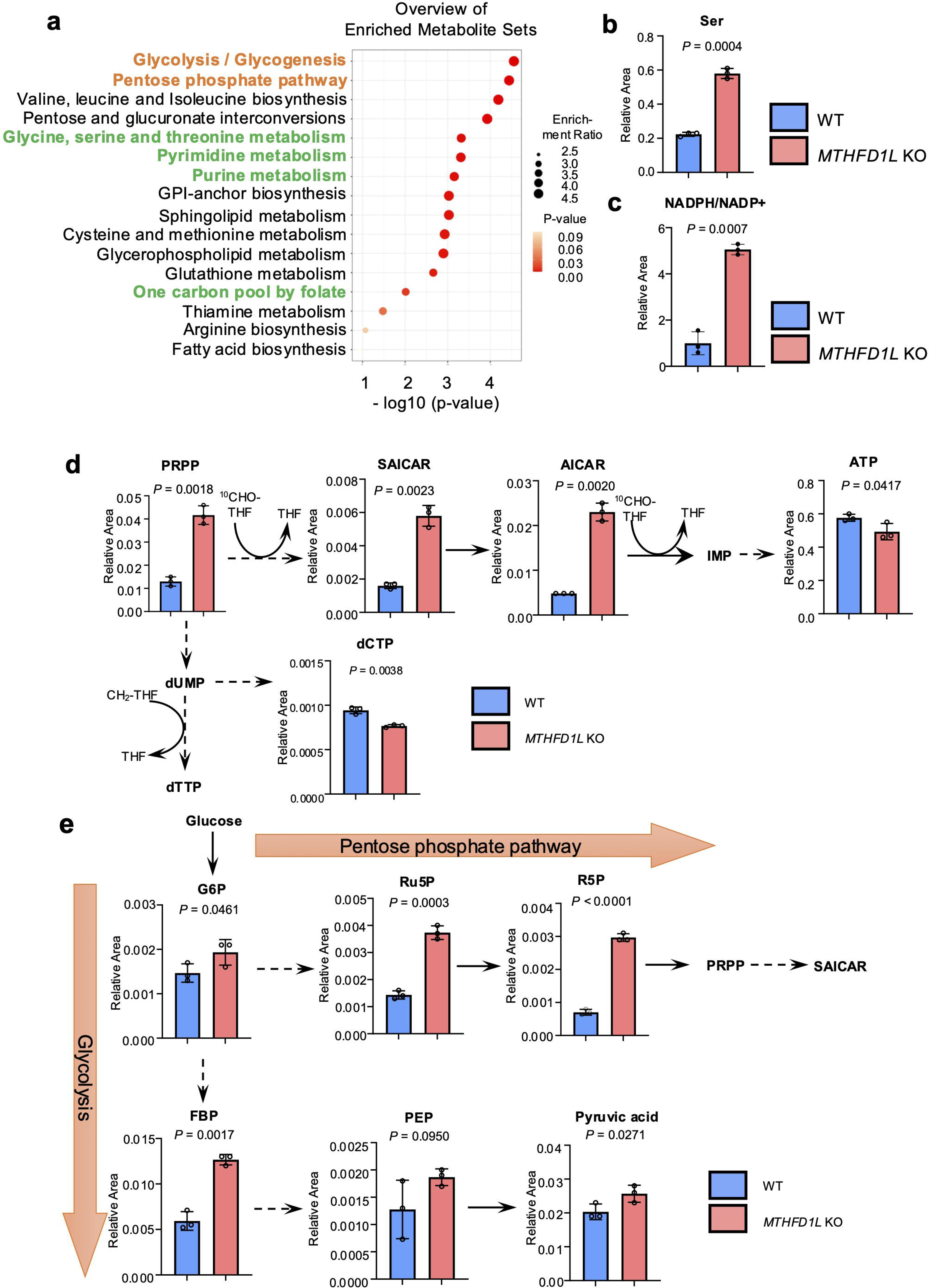
*MTHFD1L* depletion induces metabolic rewiring and accumulation of nucleotide biosynthetic intermediates. (a) Metabolite set enrichment analysis showing the altered by MTHFD1L deletion in HCC38 cells. (b) Relative levels of serine in WT and *MTHFD1L-*KO cells. (c) The NADPH/NADP^+^ ratio in WT and *MTHFD1L*-KO cells. (d) Relative abundance of metabolites involved in the pentose phosphate pathway and *de novo* nucleotide biosynthesis. *MTHFD1L* depletion resulted in accumulation of PRPP, SAICAR, and AICAR, together with alterations in nucleotide pools. (e) Relative abundance of metabolites involved in glycolysis. Dashed lines indicate multiple enzymatic reactions. Data are shown as mean ± SD. P values were determined by one-tailed Welch’s t-test. n = 3 biologically independent samples per group. PRPP (phosphoribosyl pyrophosphate), SAICAR (phosphoribosylaminoimidazolesuccinocarboxamide), AICAR (5-aminoimidazole-4-carboxamide ribonucleotide), G6P (glucose-6-phosphate), Ru5P (ribulose-5-phosphate), R5P (ribose-5-phosphate), FBP (fructose-1,6-bisphosphat), and PEP (phosphoenolpyruvate).

One-carbon units (^10^CHF-THF and CH_2_-THF) are required at several steps of de novo purine and pyrimidine synthesis (Figure 4d), and disruption of mitochondrial formate production is therefore expected to create a bottleneck within these pathways. Indeed, the purine biosynthetic intermediates PRPP, SAICAR, and AICAR accumulated markedly in MTHFD1L-KO cells (Figure 4d). Because expression levels of the enzymes responsible for generating these metabolites remained unchanged (Figure S1b), their accumulation likely reflects impaired utilization resulting from insufficient availability of one-carbon donors. Consistent with impaired nucleotide biosynthesis, ATP and dCTP levels were significantly reduced following *MTHFD1L* deletion (Figure 4d).

Unexpectedly, metabolic perturbations extended beyond nucleotide synthesis. Glycolysis and the pentose phosphate pathway were among the most significantly enriched pathways identified by metabolomic analysis (orange letters in Figure 4a). Several intermediates of these pathways, including G6P, Ru5P, R5P, FBP, PEP, and pyruvate, accumulated in MTHFD1L-KO cells (Figure 4e). RNA-seq analysis showed no substantial alterations in the expression of key glycolytic or pentose phosphate pathway enzymes, including the rate-limiting enzymes *G6PD* and *PFK* isoforms (*PFKP*, *PFKM* and *PFKL*)(Figure S2). These findings suggest that disruption of mitochondrial 1C metabolism induces widespread metabolic rewiring through accumulation of upstream intermediates and the creation of a nucleotide biosynthetic bottleneck.

Together, these results demonstrate that MTHFD1L is required to maintain metabolic homeostasis and that its loss leads to profound metabolic reprogramming characterized by accumulation of nucleotide biosynthetic intermediates and perturbation of interconnected metabolic pathways.

### 3.3. MTHFD1L is required for tumor initiation in vivo

To determine whether MTHFD1L contributes to tumorigenesis in vivo, breast cancer cells were orthotopically implanted into mammary fat pads and tumor growth was monitored. DOX-induced deletion of *MTHFD1L* significantly suppressed tumorigesis compared with control tumors (Figure 5a). Efficient Cas9 induction and *MTHFD1L* deletion in individual tumors were confirmed by immunoblot analysis (Figure 5b).

**Fig. 5.**
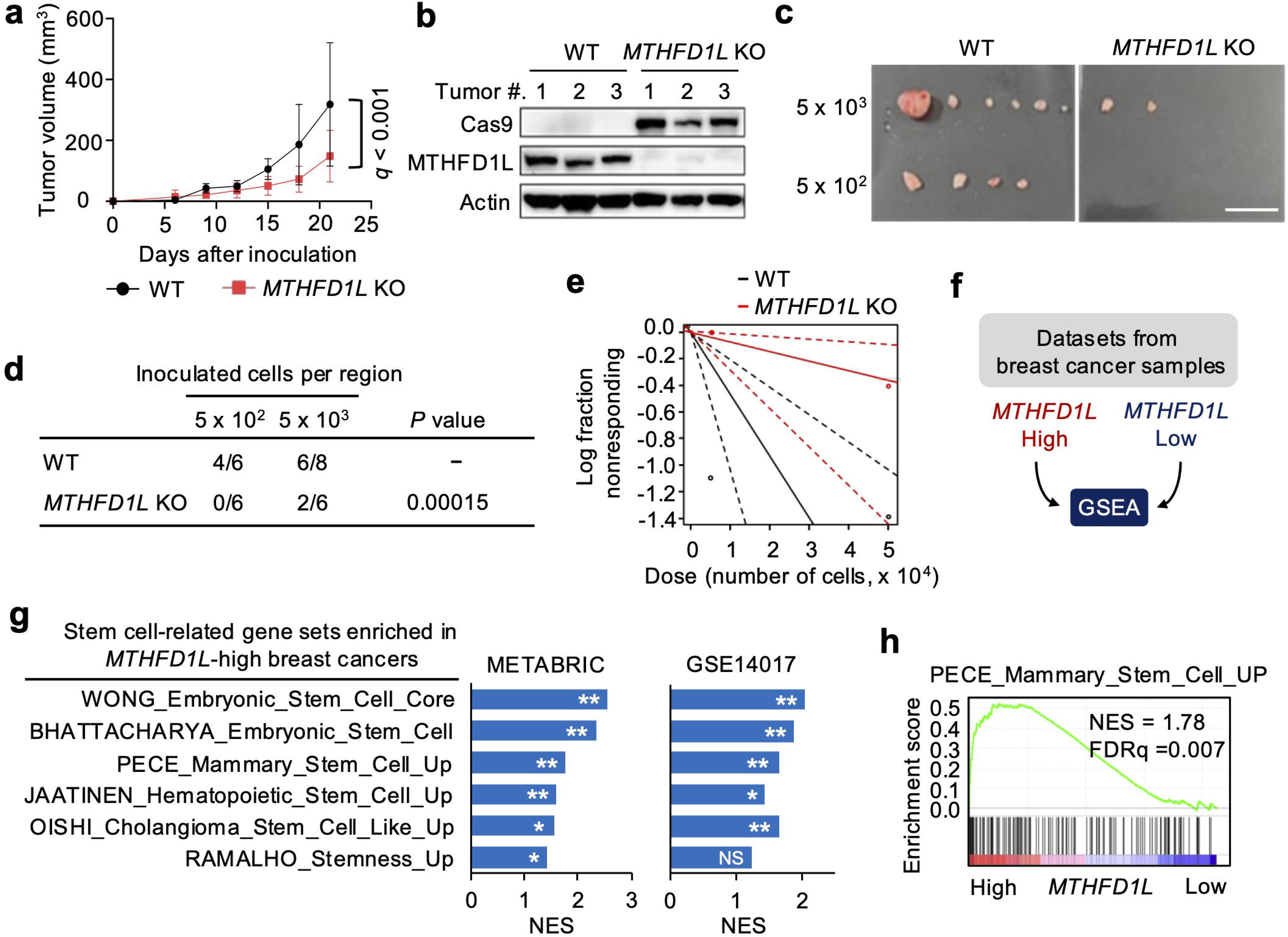
MTHFD1L is required for tumorigenesis and tumor-initiating capacity in vivo. (a) Growth of WT and *MTHFD1L*-KO HCC38 xenograft tumors. n = 5 mice for the WT group and n = 6 mice for the *MTHFD1L*-KO group. q values were determined by two-way ANOVA with the two-stage step-up method of Benjamini, Krieger, and Yekutieli. (b) Efficiency of *MTHFD1L* depletion in tumor tissues dissected from mice. Three tumors from each group were analyzed. (c–e) ELDA of WT and *MTHFD1L*-KO HCC38 cells transplanted into mice. Representative images of tumors 54 days after transplantation are shown in (c), and tumor formation frequency is shown in (d,e). Scale bar, 20 mm. P value in (d) was determined by goodness-of-fit test. (f) Schematic illustration of the gene set enrichment assay (GSEA) used to analyze breast cancer patient datasets stratified according to *MTHFD1L* expression. (g) Stem cell-associated gene sets enriched in breast cancer samples with high *MTHFD1L* expression in the METABRIC discovery dataset (left) and GSE14017 dataset (right). The C2 collection in MSigDB was used for analysis. TNBC samples from the METABRIC discovery dataset, stratified by upper and lower quantiles of *MTHFD1L* expression, and distant metastasis samples from GSE14017 were analyzed. (h) GSEA plot showing enrichment of a mammary stem cell signature in breast cancer samples from the METABRIC discovery dataset ranked according to *MTHFD1L* expression. FDR values in (g) and (h) were determined from P values calculated by random permutation test. *FDR < 0.25, **FDR < 0.05. NS; not significant, NES; normalized enrichment score.

We next evaluated tumor-initiating capacity using limiting dilution transplantation assays. *MTHFD1L* deletion substantially reduced tumor initiation frequency. At an inoculum of 5 × 10^3^ cells, tumors developed in 6/8 mice in the control group compared with 2/6 mice in the KO group, whereas at 5 × 10^2^ cells, tumors arose in 4/6 control mice but in none of the KO mice (Figure 5c–e).

To further explore the relationship between MTHFD1L and stemness, we performed GSEA using publicly available breast cancer datasets(Figure 5f)^19,20^. Stem cell-associated gene signatures were consistently enriched in tumors expressing high levels of *MTHFD1L* compared with tumors expressing low levels of *MTHFD1L* across multiple datasets (Figure 5g,h).

Collectively, these findings demonstrate that MTHFD1L is required for efficient tumor initiation and support a role for MTHFD1L in maintaining stem-like properties in breast cancer.

### 3.4. MTHFD1L is required for efficient lung colonization and maintenance of stem-like properties

Because the lung is a common site of breast cancer metastasis, we next investigated whether MTHFD1L contributes to lung colonization. Breast cancer cells were injected into the tail veins of immunodeficient NSG mice to establish experimental lung metastasis models (Figure 6a,e). Deletion of *MTHFD1L* markedly suppressed lung metastasis in both established TNBC cell lines (Figure 6b-d) and patient-derived breast cancer models (Figure 6f,g).

**Fig. 6.**
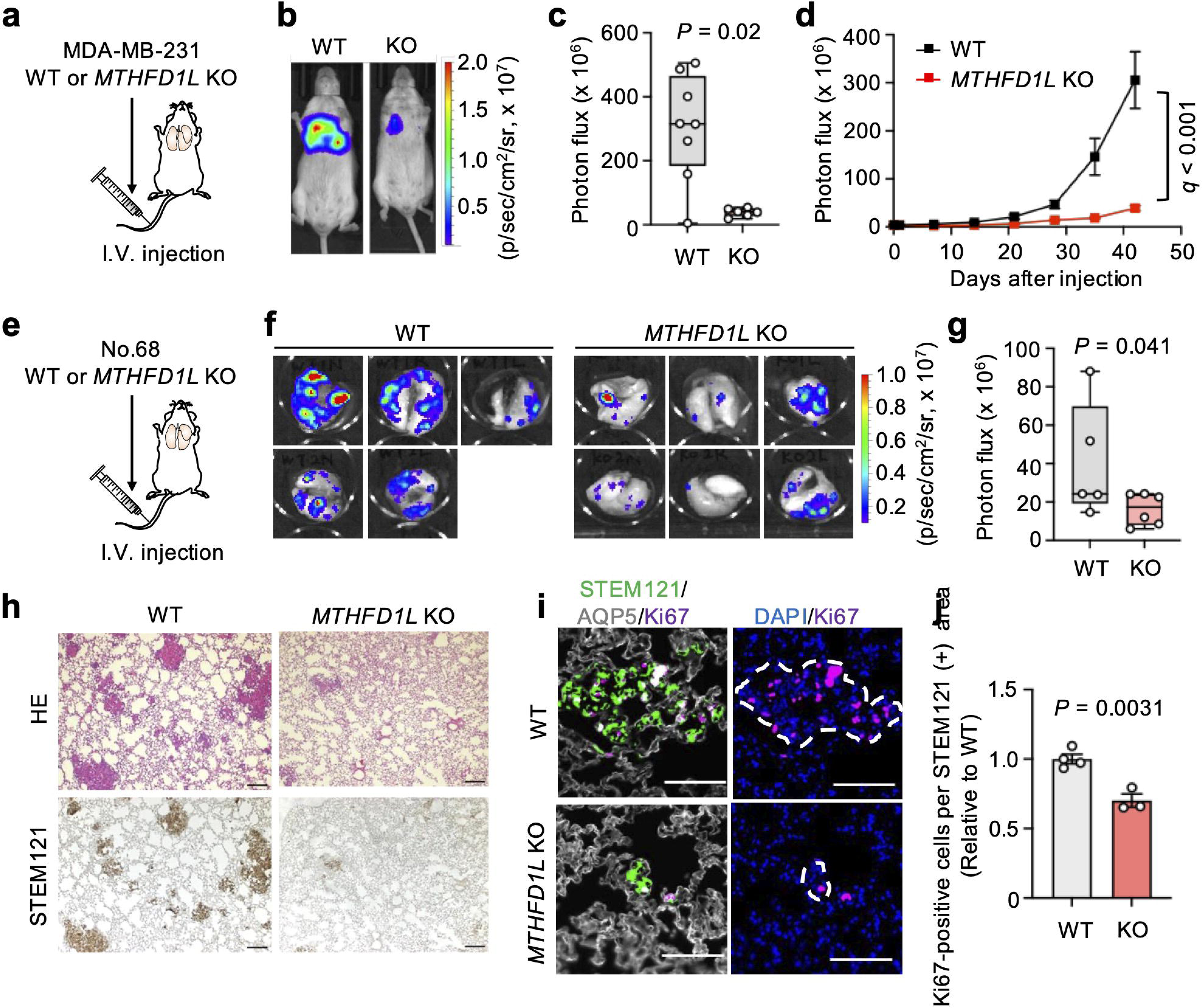
MTHFD1L is required for efficient lung colonization by breast cancer cells. (a–d) Lung colonization by WT and *MTHFD1L*-KO MDA-MB-231 cells. Schematic illustration of the experimental design (a), representative bioluminescence images (b), photon flux 42 days after intravenous injection (c), and lung colonization over time (d) are shown. I.V.; intravenous. Boxes in (c) indicate the median and interquartile range, and whiskers indicate the minimum and maximum values. P or q values were determined by two-tailed Mann–Whitney test in (c) and by two-way ANOVA with the two-stage step-up method of Benjamini, Krieger, and Yekutieli in (d). n = 8 mice for the WT group and n = 6 mice for the *MTHFD1L*-KO group. (e–g) Lung colonization by WT and *MTHFD1L*-KO No.68 patient-derived breast cancer cells. Schematic illustration of the experimental design (e), representative bioluminescence images of dissected lungs (f), and quantification of lung colonization 71 days after intravenous injection (g) are shown. P value was determined by one-tailed Mann–Whitney test. n = 5 mice for the WT group and n = 6 mice for the *MTHFD1L*-KO group. (h) Histological analysis of lung lesions by hematoxylin and eosin staining (upper panels) and immunostaining for the human-specific marker STEM121 (lower panels) in lungs of mice harboring MDA-MB-231 metastases. Scale bar, 200 µm. (i) Immunofluorescence staining for Ki67, STEM121, and aquaporin 5 (AQP5) in mouse lungs harboring MDA-MB-231 metastases. STEM121 marks human cancer cells, and AQP5 marks type I lung epithelial cells. Nuclei were stained with DAPI. Dashed lines indicate the margins of metastatic foci. Scale bar, 100 µm. (j) Quantification of Ki67-positive cells in metastatic regions. P value was determined by unpaired two-tailed t-test. n = 4 mice for the WT group and n = 3 mice for the *MTHFD1L*-KO group.

Immunohistochemical staining for the human-specific marker STEM121 confirmed a dramatic reduction in metastatic tumor burden within the lungs of mice receiving MTHFD1L-KO cells (Figure 6h). Furthermore, Ki67 staining revealed significantly reduced proliferative activity within residual metastatic lesions (Figure 6i,j).

To determine whether MTHFD1L contributes to stem-like properties during metastatic colonization, GFP-positive metastatic cancer cells were isolated from lung tissues and analyzed for stemness-associated gene expression (Figure 7a). Quantitative PCR demonstrated a significant reduction in stemness marker *SOX2* expression^14^ following *MTHFD1L* deletion (Figure 7b). Consistent with this observation, immunohistochemical analysis confirmed reduced SOX2 protein expression in metastatic lesions derived from MTHFD1L-KO cells (Figure 7c,d).

**Fig. 7.**
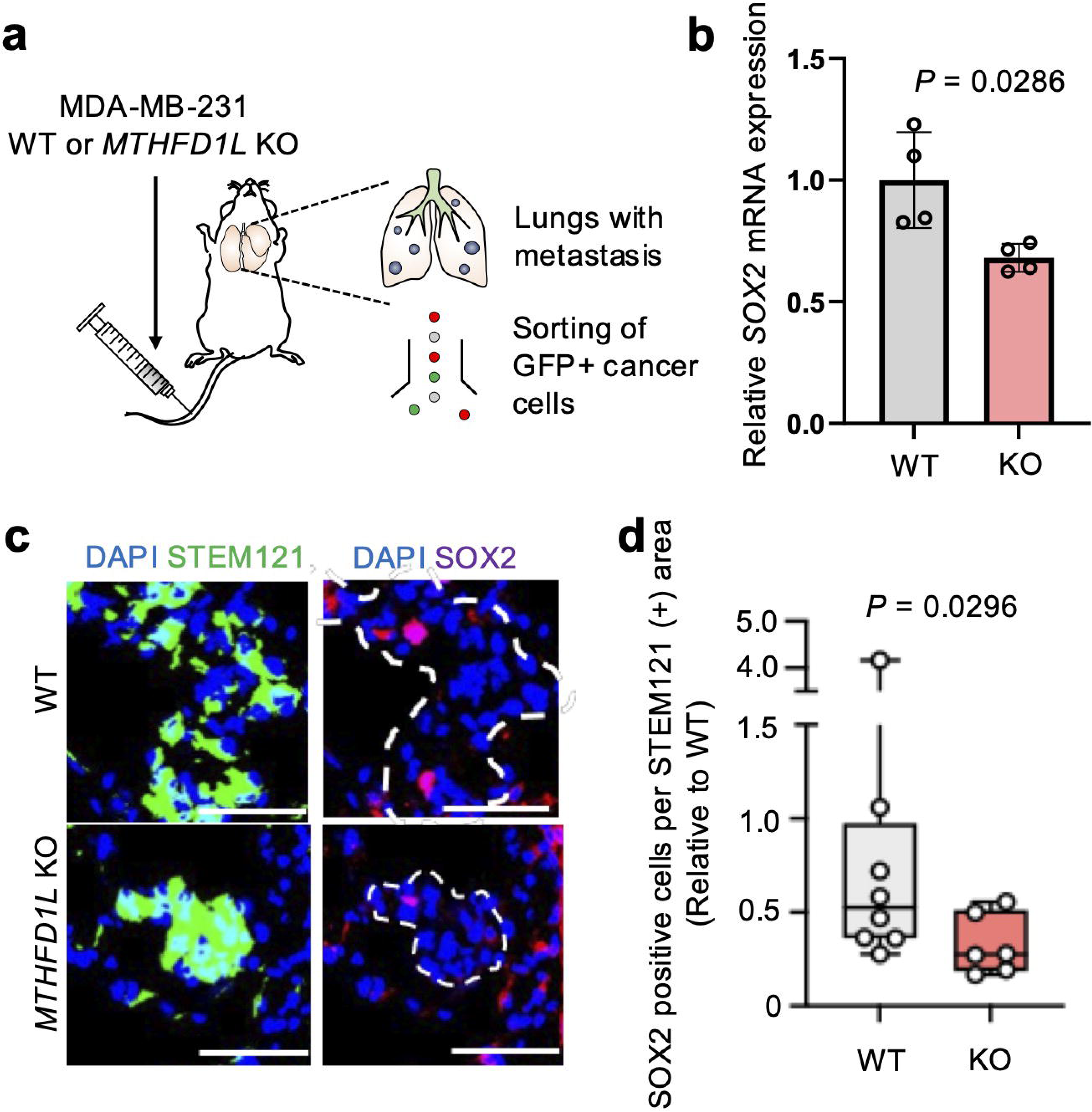
MTHFD1L sustains SOX2 expression in metastatic breast cancer cells. (a) Schematic illustration of the experimental design used to isolate metastatic breast cancer cells from lung tissues. (b) SOX2 expression in WT and *MTHFD1L*-KO MDA-MB-231 cells isolated from lungs. Data are shown as mean ± SEM. n = 4 mice for the WT group and n = 4 mice for the *MTHFD1L*-KO group. P value was determined by two-tailed Mann–Whitney test. (c) Immunofluorescence images of SOX2 in metastatic lung lesions. MDA-MB-231 cells were stained with anti-human STEM121 antibody, and nuclei were visualized with DAPI. Dashed lines indicate the margins of metastatic foci. Scale bar, 50 µm. (d) Quantification of SOX2-positive cells in metastatic foci. Boxes indicate the median and interquartile range, and whiskers indicate the minimum and maximum values. P value was determined by one-tailed Mann–Whitney test. n = 8 mice for the WT group and n = 6 mice for the *MTHFD1L*-KO group.

These findings indicate that MTHFD1L is required for efficient lung colonization and supports both proliferative capacity and stem-like properties at metastatic sites.

## 4. Discussion

In the present study, we demonstrate that the mitochondrial 1C metabolic enzyme MTHFD1L plays a critical role in maintaining stem-like properties, tumor-initiating capacity, and metastatic progression in TNBC. Using both established TNBC cell lines and patient-derived breast cancer models, we show that genetic depletion of *MTHFD1L* impairs proliferation, suppresses sphere-forming ability, reduces tumor-initiating capacity, and markedly inhibits lung colonization. These findings identify MTHFD1L as an important metabolic dependency in aggressive breast cancer and support the therapeutic potential of targeting mitochondrial 1C metabolism.

Mitochondrial 1C metabolism has emerged as an attractive therapeutic target in cancer. Among its components, SHMT2 and MTHFD2 have been extensively investigated, leading to the development of several pharmacological inhibitors, although none has yet advanced to clinical application ^21–28^. In contrast, the biological functions of MTHFD1L remain comparatively underexplored. Nevertheless, accumulating evidence suggests that MTHFD1L contributes to multiple aspects of tumor progression. For example, MTHFD1L-dependent formate production promotes tumorigenesis and invasiveness in brain tumors^29^, while MTHFD1L knockdown suppresses the enhanced migratory capacity induced by MTX treatment in cancer cells^30^. Our findings expand the current understanding of mitochondrial 1C metabolism by identifying MTHFD1L as a critical regulator of tumor-initiating capacity and metastatic colonization of the lung in breast cancer.

A notable finding of this study is the close association between MTHFD1L activity and stem-like properties. Accumulating evidence indicates that cancer stem-like cells possess distinct metabolic programs that support self-renewal, survival under stress, and metastatic competence^2^. Consistent with this concept, MTHFD1L depletion reduced sphere-formation, impaired tumor initiation in vivo, and suppressed expression of the stemness-associated transcription factor SOX2 in metastatic lesions. These observations suggest that mitochondrial 1C metabolism contributes to the maintenance of stem-like states that facilitate tumor propagation and metastatic outgrowth. Future studies will be required to determine the precise molecular mechanisms linking 1C metabolic flux to stemness-associated transcriptional programs.

Another important observation is the extensive metabolic rewiring induced by MTHFD1L depletion. Consistent with the established role of mitochondrial 1C metabolism in nucleotide biosynthesis, loss of MTHFD1L resulted in the accumulation of the purine biosynthetic intermediates PRPP, SAICAR, and AICAR, accompanied by reductions in nucleotide pools. Because expression of the enzymes responsible for generating these metabolites remained largely unchanged, their accumulation is likely attributable to impaired metabolic flux resulting from insufficient 1C unit availability. Notably, metabolic perturbations extended beyond nucleotide synthesis and affected glycolysis and the pentose phosphate pathway, highlighting the broad metabolic consequences of disrupting mitochondrial formate production. The absence of major transcriptional changes in metabolic enzymes further supports the notion that alterations in metabolic flux, rather than gene-expression reprogramming, underlie these effects.

Among the metabolites altered by MTHFD1L depletion, AICAR and SAICAR are of particular interest because of their reported roles in regulating cancer cell behavior. AICAR is a well-characterized activator of AMP-activated protein kinase (AMPK) and has been shown to suppress proliferation and stem-like properties in not only cancer cells but also normal cells ^31, 32, 33^. We previously demonstrated that MTHFD2 contributes to maintenance of stemness through regulation of AICAR levels in lung cancer cells^14^. It is therefore conceivable that accumulation of AICAR contributes, at least in part, to the suppression of stem-like properties observed following MTHFD1L depletion. In addition, SAICAR has been reported to interact with pyruvate kinase M2 (PKM2) and modulate non-canonical signaling pathways involved in cell proliferation^34–37^. Whether the marked accumulation of SAICAR observed in MTHFD1L-deficient cells represents an adaptive response that supports cancer cell survival warrants further investigation.

The metabolic rewiring induced by MTHFD1L depletion may also have important implications for cellular redox homeostasis. We observed widespread alterations in glycolysis and the pentose phosphate pathway, accompanied by an increase in the NADPH/NADP+ ratio following *MTHFD1L* depletion. Because NADPH serves as a major reducing equivalent required for antioxidant systems, including glutathione-dependent detoxification pathways, these findings suggest that redox balance is largely preserved in MTHFD1L-deficient TNBC cells despite profound perturbations in 1C metabolism. Interestingly, a previous study in hepatocellular carcinoma reported that MTHFD1L depletion decreased the NADPH/NADP+ ratio and rendered cancer cells more vulnerable to oxidative stress ^38^. The basis for these apparently contrasting observations remains unclear. One possible explanation is that compensatory NADPH-generating pathways are engaged differently across tumor types. These may include mitochondrial and cytosolic enzymes involved in folate metabolism, the pentose phosphate pathway, and other NADPH-producing reactions. More broadly, these findings suggest that the metabolic consequences of MTHFD1L inhibition are highly context-dependent and influenced by lineage-specific metabolic programs and adaptive rewiring mechanisms.

Several limitations of this study should be acknowledged. Although our findings establish an essential role for MTHFD1L in TNBC growth and lung colonization, the downstream mechanisms linking 1C metabolism to stemness remain incompletely understood. In addition, the present study relied primarily on genetic depletion approaches, and future studies employing pharmacological inhibition of MTHFD1L will be important for evaluating its therapeutic tractability. Further investigation will also be required to determine whether the dependency on MTHFD1L extends to additional breast cancer subtypes and other malignancies. Nevertheless, our findings support a model in which MTHFD1L functions as a central node connecting 1C metabolism, metabolic plasticity, and stem-like phenotypes in aggressive breast cancer.

In summary, our study identifies MTHFD1L as a previously underappreciated regulator of stem-like properties, tumor initiation, and lung colonization in breast cancer. MTHFD1L functions as a central node linking mitochondrial 1C metabolism to metabolic rewiring, redox homeostasis, and cancer stem-like phenotypes. By coordinating these interconnected processes, MTHFD1L promotes tumor initiation and metastatic colonization, providing a strong rationale for the development of therapeutic strategies targeting MTHFD1L and related mitochondrial 1C metabolic pathways in aggressive breast cancer.

## Acknowledgement

A lentiviral vector encoding a triple-reporter fusion gene (TGL) consisting of herpes simplex virus thymidine kinase (HSV1-TK), green fluorescent protein (GFP), and firefly luciferase (Fluc). The expression vector was kindly provided by Dr. Thordur Oskarsson (Department of Molecular Oncology, H. Lee Moffitt Cancer Center & Research Institute, Tampa, Florida, USA).

## Funding Information

This work was supported by WISE Program of Kanazawa University by MEXT to H. Kusunoki. This work was supported in part by The Uehara Memorial Foundation, Kobayashi Foundation for Cancer Research and Kieiotoyo Foundation to T. Hongu. This work was supported in part by JSPS KAKENHI Grant Numbers JP23K18237, JP24K02304, and JP25K22575, a research grant from AMED Project for Cancer Research and Therapeutic Evolution (21447913, 21446781, 24028177, 24021759), and the Uehara Memorial Foundation, the Princess Takamatsu Cancer Research Fund, the Takeda Science Foundation, the Chugai Foundation for Life Science, and the Yasuda Medical Foundation to N. Gotoh. This work was supported in part by MEXT Promotion of Development of a Joint Usage/Research System Project: Coalition of Universities for Research Excellence Program (CURE), Grant Number JPMXP1323015484 to T. Hongu and N. Gotoh.

## Conflicts of Interest

Noriko Gotoh is Associate Editor for Cancer Science.

## Ethics Statement

- Approval of the research protocol by an Institutional Reviewer Board

The use of the patient samples was approved by the Institutional Review Boards of the Cancer Research Institute of Kanazawa University, Kanazawa University Hospital, the University of Tokyo Hospital (approval no. 72331-20).

- Informed Consent: N/A

- Registry and the Registration No. of the study/trial: N/A

- Animal Studies: All animal studies were approved by the Animal Research Committee of Kanazawa University.

## Notes

### Competing Interest Statement

The authors have declared no competing interest.

