## Supplementary information for "Mitochondrial one-carbon enzyme MTHFD1L sustains stemness and metastatic progression in breast cancer"

#### Supplementary materials and methods

##### Cell culture

MDA-MB-231, BT20, HCC1143, HCC38, MDA-MB-436 and HEK293T cells were obtained from ATCC. MDA-MB-231 and HEK293T cells were cultured in Dulbecco's Modified Eagle Medium (DMEM) (Nacalai Tesque, Kyoto, Japan) supplemented with 10% fetal bovine serum (FBS, Thermo Fisher Scientific, Waltham, MA). BT20, HCC1143, HCC38 and MDA-MB-436 cells were cultured in Roswell Park Memorial Institute 1640 (RPMI-1640) (Nacalai Tesque, Kyoto, Japan) supplemented with 10% FBS.

Patient-derived breast cancer cells (No.68, K24, K53, and K66) were established from triple negative subtype breast cancer specimens obtained from the University of Tokyo Hospital or Kanazawa University Hospital (Table S1) [1,2]. The use of these samples was approved by the Institutional Review Boards of the Cancer Research Institute of Kanazawa University, Kanazawa University Hospital, the University of Tokyo Hospital (approval no. 72331-20). These cells were maintained in organoid medium prepared from Advanced DMEM/F12 (Gibco) supplemented with the components listed (Table S2). Hydrocortisone (0.48 μg/mL; STEMCELL Technologies, Vancouver, British Columbia, Canada) was included in the culture medium. For experiments using doxycycline (DOX)-inducible systems, Tet-system-approved FBS (Takara Bio, Kyoto, Japan) was used. All media contained 50 U/ml penicillin and 50 μg/ml streptomycin (Nacalai Tesque). All cells were cultured at 37°C in a humidified atmosphere containing 5% CO2.

##### Sphere formation assay

Sphere formation assays were performed as previously described [1,2]. Cells were plated in ultra-low attachment 24-well plates (Corning) at a density of 2,000 cells per well and cultured in sphere culture medium consisting of DMEM/F12 supplemented with 20 ng/mL epidermal growth factor (EGF) (Merck Millipore, Burlington, MA), 20 ng/mL basic fibroblast growth factor (bFGF)(PeproTech, Cranbury, NJ), B27 supplement (Thermo Fisher Scientific), and 4 μg/mL heparin (STEMCELL Technologies). After 7 days, spheres with diameters greater than 75 μm were counted.

##### In vitro limiting dilution assay

Single-cell suspensions were prepared using Accumax (Innovative Cell Technologies). Cells were seeded into ultra-low attachment 96-well plates at serial dilutions ranging from

200 to 6 cells per well. After culture, wells containing spheres larger than 75 μm were scored as positive. Sphere-forming frequency was calculated using Extreme Limiting Dilution Analysis (ELDA) software (https://bioinf.wehi.edu.au/software/elda/).

#### Analysis of gene expression profiles

*MTHFD1L* expression according to breast cancer grade and molecular subtype was analyzed using the METABRIC dataset[3] through cBioPortal[4]. *MTHFD1L* expression in normal breast tissues and breast tumors was analyzed using the GSE3744[5] and TCGA-BRCA datasets. GSE3744 data were RMA-normalized using R software (version 4.5.2). Gene set enrichment analysis (GSEA) was performed using the Molecular Signatures Database (MSigDB). Survival analyses were conducted using the Kaplan–Meier Plotter database[6]. The datasets included in the Kaplan–Meier Plotter analysis are follows: GSE1456, GSE16446, GSE16716, GSE20271, GSE20685, GSE20711, GSE22093, GSE3494, GSE37946, GSE42568, GSE45255, GSE48390, GSE58812, GSE65194, GSE69031, GSE7390 for analysis of overall survival; GSE11121, GSE16446, GSE16716, GSE17907, GSE19615, GSE20685, GSE22093, GSE25066, GSE26971, GSE2990, GSE3494, GSE45255, GSE5327, GSE58812, GSE61304, GSE65194, GSE6532, GSE69031, GSE7390, GSE9195 for distant metastasis free survival.

#### Lentivirus-mediated knockdown

shRNAs targeting human *MTHFD1L* were cloned into lentiviral vectors and transfected into HEK293T cells using Lipofectamine 2000 (Thermo Fisher Scientific). Viral supernatants were collected 48 h after transfection and used to infect BT20 cells in the presence of 8 μg/mL polybrene (Nacalai Tesque). Infected cells were selected with puromycin (Nacalai Tesque) (2 μg/mL). Knockdown efficiency was confirmed by quantitative PCR and western blotting. The shRNA sequences used in this study are listed in Table S3.

#### CRISPR/Cas9-mediated knockout

DOX-inducible Cas9 expressing cells were generated as described[7]. Briefly, Edit-R-inducible lentiviral Cas9 nuclease (Horizon Discovery, Cambridge, UK) and lentiviral packaging plasmids (pCMV-VSV-G-RSV-Rev and pCAG-HIVgp, kindly provided by Dr. H. Miyoshi [RIKEN]) were co-transfected onto lentiX-293T (Takara Bio, USA) or HEK293T cells using PLUS reagent and lipofectamine or lipofectamine 2000 (Thermo Fisher Scientific) according to the manufacturer's instructions. Supernatant containing lentivirus was added to breast cancer cells attached to a culture dish. Infected cells were selected in medium supplemented with 3 µg/mL blasticidin S HCl (Thermo Fisher Scientific).

A plasmid expressing *MTHFD1L*-specific sgRNA was constructed by using the pLenti-sgRNA (Addgene #71409). The virus for expressing sgRNA was prepared and added to the culture of DOX-inducible Cas9 expressing cells. Two days after infection, infected cells were selected in medium containing 2.5 µg/mL puromycin (Nacalai Tesque). Subsequently, single-cell clones were established. The efficiency of Cas9 expression and MTHFD1L KO was evaluated by western blotting after culturing cells in medium supplemented with 1 µg/ml DOX (Tokyo Chemical Industries, Tokyo, Japan). The target sequences of the sgRNAs for *MTHFD1L* used are as follows: 5'-GAATTTGGCTGAGGAGGTGA-3'.

**Cell growth assay**

Cells were seeded in 96-well plates at densities of 500–1,000 cells per well depending on the cell line. After 5 days of culture, cell viability was assessed using the CellTiter-Glo Luminescent Cell Viability Assay (Promega, Madison, WI) according to the manufacturer's instructions. Luminescence signals were measured using a microplate reader and normalized to control samples.

**Metabolome analysis**

HCC38 cells were washed twice with 3.4% meso-erythritol, and intracellular metabolites were extracted using methanol containing internal standards (Human Metabolome Technologies [HMT], Tsuruoka, Japan). The extracts were collected and centrifuged through a 5-kDa cutoff ultrafiltration filter (HMT) at 9,100 × g for 2–5 h at 4°C to remove macromolecules.

Metabolome analysis was conducted according to HMT's *Basic Scan* package, using capillary electrophoresis time-of-flight mass spectrometry (CE-TOFMS) based on the methods described previously [8,9]. Briefly, CE-TOFMS analysis was carried out using an Agilent CE capillary electrophoresis system equipped with an Agilent 6230 time-of-flight mass spectrometer (Agilent Technologies, Santa Clara, CA). The systems were controlled by Agilent MassHunter Workstation Data Acquisition (Agilent Technologies) and connected by a fused silica capillary (50 µm *i.d.* × 62 cm total length) with commercial electrophoresis buffer (H3301-1001 and I3302-1023 for cation and anion analyses, respectively, HMT) as the electrolyte. The spectrometer was scanned from *m/z* 50 to 1,000 and peaks were extracted using MasterHands, automatic integration software (Keio University, Tsuruoka, Japan) in order to obtain peak information including *m/z*, peak area, and migration time (MT)[10]. Signal peaks corresponding to isotopomers, adduct ions, and other product ions of known metabolites were excluded, and the remaining peaks were annotated according to HMT's metabolite database based on their *m*/*z* values and MTs. Areas of the annotated peaks were then normalized to internal standards and sample amount in order to obtain relative levels of each metabolite. Primary 110 metabolites were absolutely quantified based on one-point calibrations using their respective standard compounds. Enrichment analysis was performed using MetaboAanalyst 6.0[11].

**Generation of luciferase-expressing breast cancer cells**

For in vivo lung colonization assays, MDA-MB-231 and No.68 cells were transduced with a lentiviral vector encoding a triple-reporter fusion gene (TGL) consisting of herpes simplex virus thymidine kinase (HSV1-TK), green fluorescent protein (GFP), and firefly luciferase (Fluc)[12]. Lentiviral particles were produced in HEK293T cells by co-transfection of the TGL vector with pMD2.G and psPAX2 packaging plasmids using Lipofectamine 2000 (Thermo Fisher Scientific). Viral supernatants were collected 48 h after transfection and used to infect target cells. GFP-positive cells were isolated by fluorescence-activated cell sorting (FACSAria III, BD Biosciences, San Jose, CA) and used for subsequent experiments.

**Tumor growth assay**

Female NOD/SCID/IL2R$\gamma^{null}$ (NSG) or SCID Beige mice aged 6–12 weeks were purchased from Ninox Lab Supply Inc. (Kanazawa, Japan) and maintained under pathogen-free conditions in accordance with institutional guidelines. All animal studies were approved by the Animal Research Committee of Kanazawa University (approval number: AP24-021-03 and AP24-023-03).

For xenograft studies, $1 \times 10^5$ HCC38 cells expressing the indicated constructs were suspended in phosphate-buffered saline (PBS) and injected into the mammary fat pads of female SCID Beige mice. Tumor volume was calculated using the formula V = 4/3 ×

$\pi \times (S/2) \times (L/2)^2$, where S and L represent the minor and major tumor diameters, respectively. For in vivo limiting dilution assays, 500 or 5,000 MDA-MB-231 cells were suspended in a 1:1 mixture of Matrigel (Corning, New York, NY) and PBS and injected into the mammary fat pads of female NSG mice. Tumor-initiating frequency was calculated using ELDA software (https://bioinf.wehi.edu.au/software/elda/).

**Flow cytometry and isolation of metastatic cancer cells**

For isolation of metastatic cancer cells from mouse lungs, lung tissues were dissociated in digestion buffer containing 0.5% collagenase type III (Pan Biotech, AidenBach, Germany), 1% dispase II (Thermo Fisher Scientific), and 30 µg/mL DNase I (NIPPON GENE, Tokyo, Japan) at 37°C for 45 min. Cell suspensions were filtered through 100-µm and 70-µm cell strainers and treated with red blood cell lysis buffer (Roche).

Cells were resuspended in fluorescence-activated cell sorting (FACS) buffer consisting of PBS supplemented with 2% FBS and 2 mM EDTA (Nakarai tesque). Dead cells were excluded using 4',6-diamidino-2-phenylindole (DAPI, 1:2000 dilution, DOJINDO, Kumamoto, Japan) staining. GFP-positive and DAPI-negative cells were sorted using a FACSAria III cell sorter (BD Biosciences) and used for downstream analyses.

**Immunofluorescence**

Mouse lungs were fixed in 4% paraformaldehyde, cryoprotected in 30% sucrose, embedded in O.C.T. compound (Sakura Finetek, Tokyo, Japan), and sectioned at 8 µm thickness. Tissue sections were blocked and incubated with primary antibodies against STEM121 (1:1000 dilution, Takara, Kyoto, Japan), SOX2(D6D9, 1:100 dilution, Cell Signaling Technology, Danver, MA), Ki67 (SolA15, 1:200 dilution, Invitrogen), or AQP5 (ab305303, 1:100 dilution, abcam, Cambridge, UK). Following incubation with fluorescence-conjugated secondary antibodies, nuclei were counterstained with DAPI.

Images were acquired using a BZ-X800 fluorescence microscope (Keyence, Osaka, Japan) and analyzed using FIJI (ImageJ).

**Immunohistochemistry**

For immunohistochemical analyses, lungs containing metastatic lesions were fixed in 4% paraformaldehyde, embedded in paraffin, and sectioned at 4–5 µm thickness as described[2]. After deparaffinization and antigen retrieval (pH9.0), sections were blocked and incubated with anti-STEM121 antibody (1:1000 dilution, Takara, Kyoto, Japan) overnight at 4°C.

Signal detection was performed using a horseradish peroxidase-conjugated secondary antibody and DAB substrate (NICHIREI, Tokyo, Japan) according to the manufacturers' instructions. Hematoxylin and eosin staining was performed using standard procedures. Images were acquired using a BZ-X800 microscope (Keyence, Osaka, Japan).

**RNA sequencing**

Total RNA was isolated from HCC38 cells using the RNeasy Mini Kit (Qiagen, Venlo, The Netherlands). Sequencing libraries were prepared using the NEBNext Ultra II Directional RNA Library Prep Kit (New England Biolabs, Ipswich, MA), and paired-end sequencing (150 bp) was performed on a NovaSeq X Plus platform (Illumina, San Diego, CA).

Sequencing reads were trimmed using Trim Galore and aligned to the human reference genome (hg38) using STAR[13]. Gene-level read counts were generated using featureCounts[14], and differential expression analysis was performed using DESeq2[15]. RNA-seq data generated in this study have been deposited in the Gene Expression Omnibus (GEO) under accession number GSE333926.

**Western blotting**

Whole-cell lysates were prepared using standard method as described[16]. PVDF membranes were incubated with primary antibodies against MTHFD1L (#14998S, Cell Signaling Technology), Cas9 (#61577, ACTIVE MOTIF), MTHFD2 (4G7-2G3, Abnova), SHMT2 (HPA020549, Atlas Antibodies), GLDC (Abcam), or β-actin (MAB150, Merck Millipore), followed by horseradish peroxidase-conjugated secondary antibodies (Millipore).

Protein signals were visualized using chemiluminescence reagents and detected using a FUSION-FX7 imaging system (Vilber Bio Imaging).

**Quantitative RT-PCR**

Total RNA was isolated using the RNeasy Mini Kit (Qiagen), and cDNA was synthesized using the High-Capacity cDNA Reverse Transcription Kit (Applied Biosystems, Waltham,

MA). Quantitative PCR was performed using SYBR Green chemistry on a QuantStudio1 or ViiA7 Real-Time PCR System (Applied Biosystems). Relative gene expression was calculated using the ΔΔCt method. Primer sequences are listed in Table S4.

13 Dobin A, Davis CA, Schlesinger F, et al. STAR: ultrafast universal RNA-seq aligner.

*Bioinformatics*. 2013; 29: 15-21.

14 Liao Y, Smyth GK, Shi W. featureCounts: an efficient general purpose program for assigning sequence reads to genomic features. *Bioinformatics*. 2014; 30: 923-930.

15 Love MI, Huber W, Anders S. Moderated estimation of fold change and dispersion for RNA-seq data with DESeq2. *Genome Biol*. 2014; 15: 550.

16 Nishimura T, Nakata A, Chen X, et al. Cancer stem-like properties and gefitinib resistance are dependent on purine synthetic metabolism mediated by the mitochondrial enzyme MTHFD2. *Oncogene*. 2019; 38: 2464-2481.

### Figure S1

a

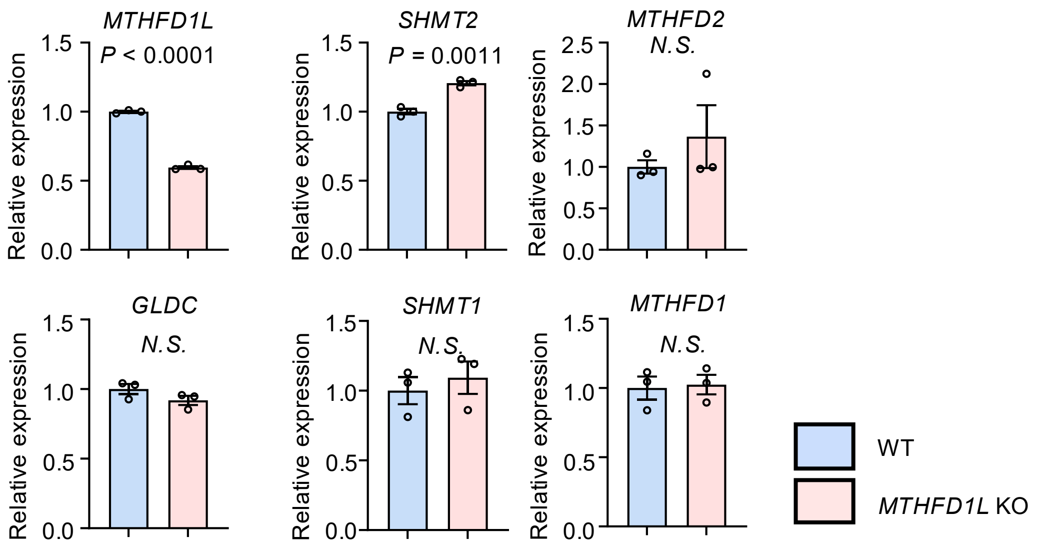

MTHFD1L
P < 0.0001
SHMT2
P = 0.0011
MTHFD2
N.S.
GLDC
N.S.
SHMT1
N.S.
MTHFD1
N.S.
Relative expression
1.5
1.0
0.5
0.0
2.5
2.0
WT
MTHFD1L KO


b

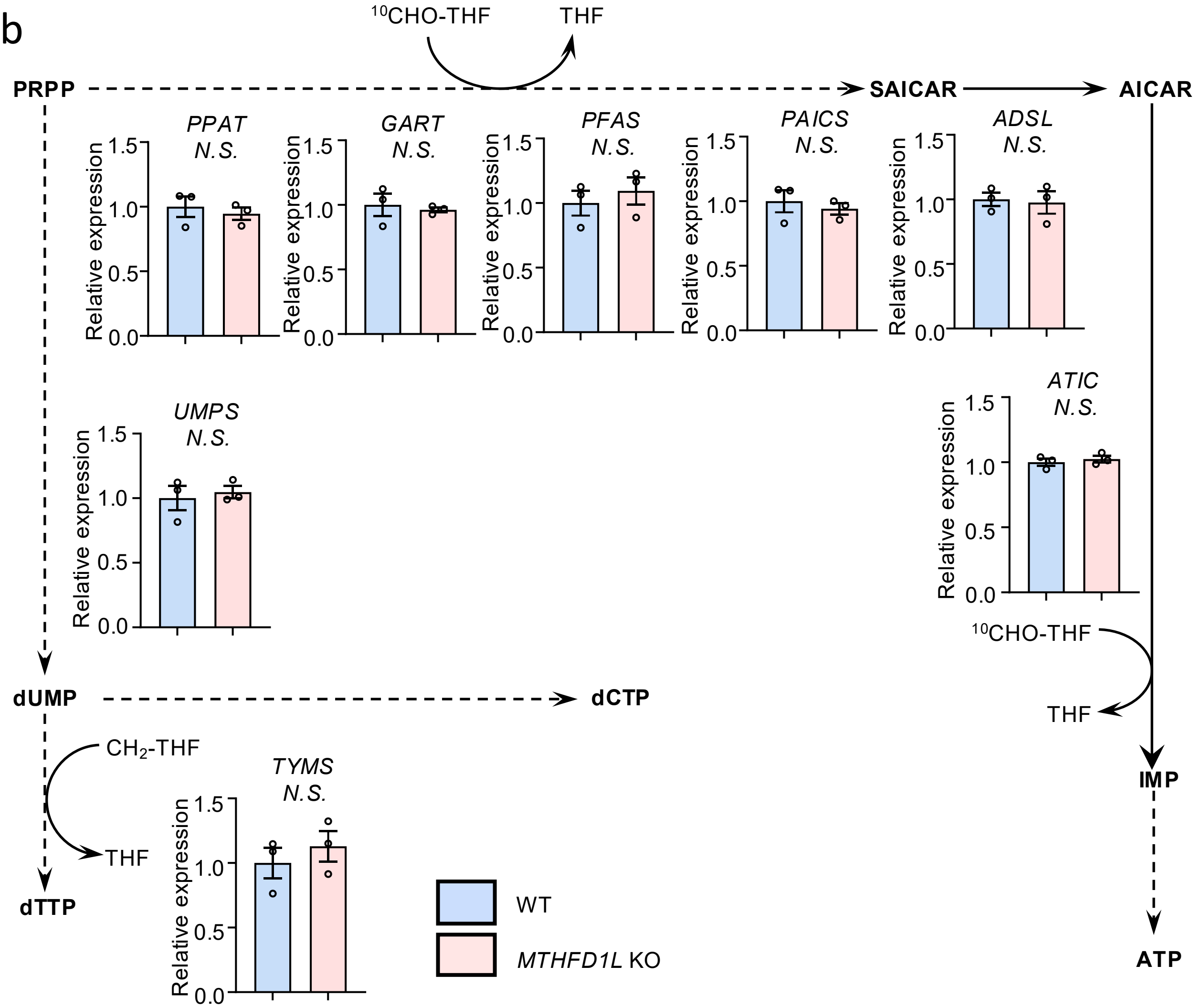

10CHO-THF
THF
PRPP
SAICAR
AICAR
PPAT
N.S.
GART
N.S.
PFAS
N.S.
PAICS
N.S.
ADSL
N.S.
ATIC
N.S.
UMPS
N.S.
TYMS
N.S.
Relative expression
1.5
1.0
0.5
0.0
dUMP
dCTP
CH2-THF
THF
dTTP
10CHO-THF
THF
IMP
ATP
WT
MTHFD1L KO

### Figure S2

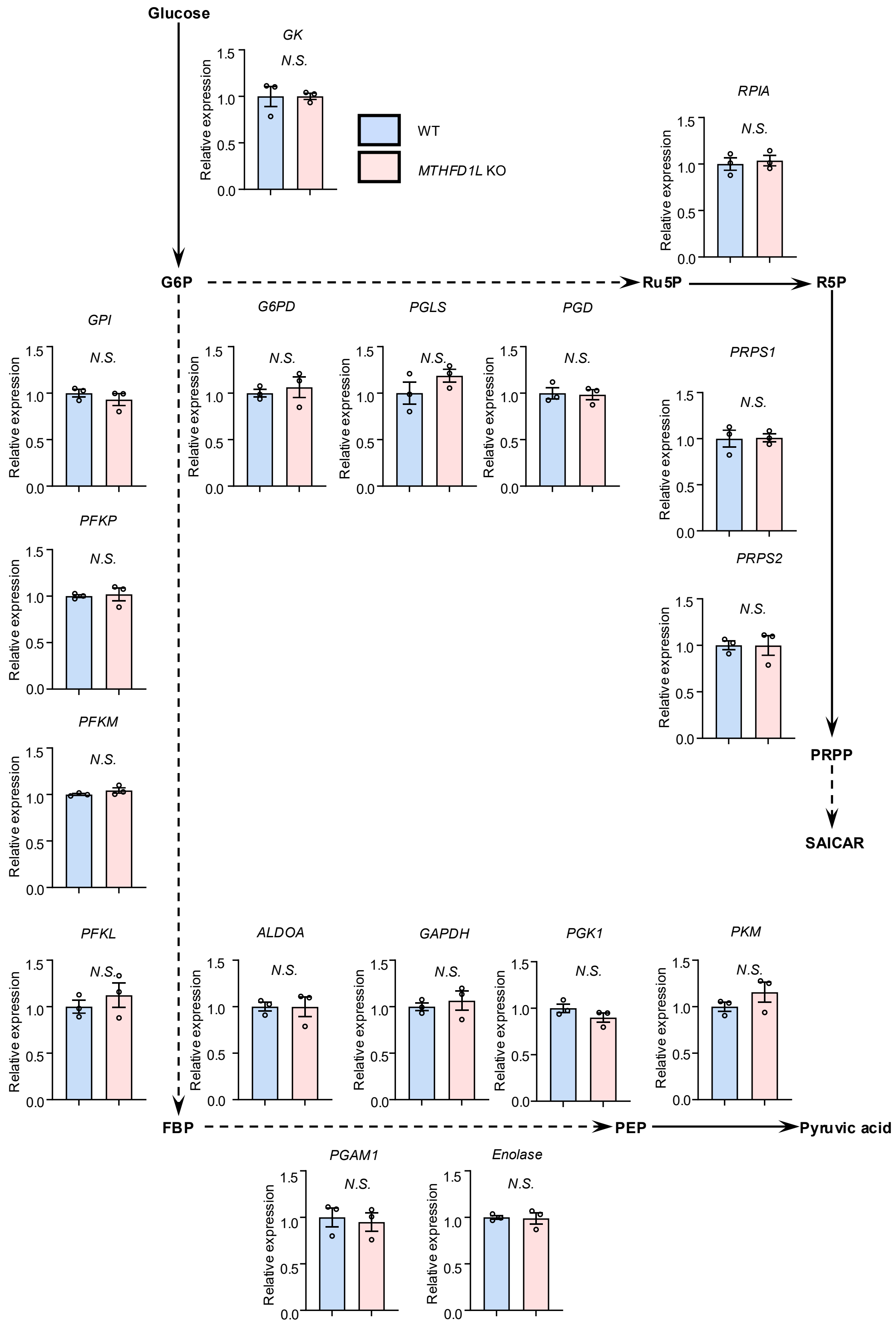

Glucose
G6P
Ru5P
R5P
PRPP
SAICAR
FBP
PEP
Pyruvic acid
WT
MTHFD1L KO
GK
RPIA
GPI
G6PD
PGLS
PGD
PRPS1
PRPS2
PFKP
PFKM
PFKL
ALDOA
GAPDH
PGK1
PKM
PGAM1
Enolase
N.S.
Relative expression
0.0
0.5
1.0
1.5

### Supplementary Table 1

Patient characteristics of breast cancer patients

| Patient | Identifer (Patient age) | Histological subtype | ER (IHC) | PR (IHC) | HER2 (IHC) | Ki67 index | Clinical subtype | BRCA1, 2 mutations | Clinical drug resistance | Stage |
|---|---|---|---|---|---|---|---|---|---|---|
| P1 | No.68 (55) | ILC | - | - | - | 50% | TNBC | - | No | IIB |
| P2 | K24 (77) | IDC | - | - | - | 80% | TNBC | BRCA1, 2 | No | IIIC |
| P3 | K53 (84) | IDC | - | - | - | | TNBC | BRCA1, 2 | No | IIA |
| P4 | K66 (57) | IDC | - | - | - | 17% | TNBC | BRCA2 | Yes | IV |

ER, estrogen receptor; PR, progesterone receptor; IHC, immunohistochemistry; IDC, invasive ductal carcinoma; ILC, invasive lobular carcinoma; ND, not determined. Patients 4 is relapsed cases after chemotherapy.

### Supplementary Table 2

Reagents used in organoid medium

| Reagents | Working Concentration |
|---|---|
| hydrocortisone (Stemcell Technology) | 1 μM |
| A83-01 (Tocris) | 500 nM |
| FGF10 (Wako) | 20 ng/mL |
| FGF7 (Wako) | 5 ng/mL |
| Neuregulin-1 (Peprotech) | 5 nM |
| EGF (Millipore) | 5 ng/mL |
| Y-27632 (Wako) | 5 μM |
| SB202190 (CAYMAN CHEMICAL COMPANY) | 500 nM |
| R-spondin3 (Peprotech) | 250 ng/mL |
| heparin (Stemcell Technology) | 4 μg/mL |
| Noggin (Wak) | 100 ng/mL |
| Primocin (invivogen) | 50 μg/mL |
| N-Acetylcysteine (Sigma) | 1.25 mM |
| Nicotinamid (Sigma) | 5 mM |
| B27 (Thermo Fisher Scientific) | x1 |
| GlutaMax (Thermo Fisher Scientific) | x1 |
| Hepes (Nacalai Tesque) | 10 mM |
| P/S (Nacalai Tesque) | x1 |

### Supplementary Table 3

shRNA oligos used for lentivirus-mediated gene knockdown.

| shRNA | Sequence |
|---|---|
| Control shRNA | 5’-TGCTGTTGACAGTGAGCGTAGATAAGCATTATAATTCCTTAGTGAAGCCA CAGATGTAAGGAATTATAATGCTTATCTACTGCCTCGGA-3’ |
| hMTHFD1L shRNA#1 | 5’-TGCTGTTGACAGTGAGCGCTCAAAAGAAGTTCTAAGTTTATAGTGAAGCC ACAGATGTATAAACTTAGAACTTCTTTTGAATGCCTACTGCCTCGGA-3’ |
| hMTHFD1L shRNA#2 | 5’-TGCTGTTGACAGTGAGCGCTCACCTGTTGCCAAAGCTGTATAGTGAAGCC ACAGATGTATACAGCTTTGGCAACAGGTGAATGCCTACTGCCTCGGA-3’ |

### Supplementary Table 4

Primers used for qPCR.

| Gene | Forward | Reverse |
|---|---|---|
| human *SOX2* | 5’-GGGGGAATGGACCTTGTATAG-3’ | 5’-GCAAAGCTCCTACCGTACCA-3’ |
| human *RPL13A* | 5’-AGATGGCGGAGGTGCAG-3’ | 5’-GGCCCAGCAGTACCTGTTTA-3’ |
